# New mining concessions will severely decrease biodiversity and ecosystem services in Ecuador

**DOI:** 10.1101/251538

**Authors:** Bitty Roy, Martin Zorrilla, Lorena Endara, Dan Thomas, Roo Vandegrift, Jesse M. Rubenstein, Tobias Policha, Blanca Rios-Touma

**Affiliations:** Institute of Ecology and Evolution, University of Oregon, Eugene, OR 97403; Nutrition Technologies Ltd./Research Institute of Biotechnology and Environment, Ho Chi Minh City, Vietnam; Department of Biology, University of Florida, Gainesville, FL 32611; Department of Geography, University at Buffalo, Buffalo, NY 14261; Grupo de Investigación en Biodiversidad Medio Ambiente y Salud ‐BIOMAS‐ Facultad de Ingenierías y Ciencias Agropecuarias, Universidad de Las Américas, Quito-Ecuador.

**Keywords:** ABVP, birds, bosque protector, cloud forest, corridors, copper mining, Ecuador, elevational gradient, endangered, frogs, jaguar, orchids, primates, protected areas, SNAP

## Abstract

Ecuador has the world’s highest biodiversity, despite being a tiny fraction of the world’s land area. The threat of extinction for much of this biodiversity has dramatically increased since April 2016, during which time the Ecuadorian government has opened approximately 2.9 million hectares of land for mining exploration, with many of the concessions in previously protected forests. Herein, we describe the system of protected lands in Ecuador, their mining laws, and outline the scale of threat by comparing the mammals, amphibians, reptiles, birds, and orchids from several now threatened protected areas, classed as “Bosques Protectores” (BPs), in the NW montane cloud forests. We examine two large (>5,000 ha) BPs, Los Cedros and El Chontal, and two medium BPs, Mashpi (1,178 ha) and Maquipucuna (2,474 ha). Since BP El Chontal is so poorly explored, we used several other small reserves (<500 hectares) in the Intag Valley to gain an idea of its biodiversity. Together, these BPs and reserves form a buffer and a southern corridor for the still-protected Cotacachi-Cayapas Ecological Reserve, which is otherwise now surrounded by mining concessions. We gathered published literature, “gray literature”, information from reserve records and websites, and our previously unpublished observations to make comparative species tables for each reserve. Our results from these still incompletely known reserves reveal the astonishing losses that mining will incur: eight critically endangered species, including two primates (brown-headed spider monkey and white-fronted capuchin), 37 endangered species, 149 vulnerable and 85 near threatened and a large number of less threatened species Our data show that each of the reserves protects a unique subset of taxa in this land of highly localized endemics. Each of the reserves also generates sustainable income for the local people. The short-term national profits from mining will not compensate for the permanent biodiversity losses, and the long-term ecosystem service and economic losses at the local and regional level.

## INTRODUCTION

### New mining concessions in Ecuador

During the years of 2016 and 2017, the Ecuadorian Ministry of Mining increased exploratory mining concessions across the country from roughly 3% to more than 13% of the country’s continental land area (Vandegrift et al., 2017). These new concessions significantly decrease forest protected areas, with more than 30% of the total land area protected by Bosques Protectores included in new exploratory mining concessions (Vandegrift et al., 2017). The majority of the concessions are located in the hyper-diverse Andean Forest Zone, composed of montane and cloud forests (Fig. 1A), the eco-region with the highest biodiversity in the region (Gentry, 1992)and one of the most threatened eco-regions on the planet (Myers, Mittermeier, Mittermeier, da Fonseca, & Kent, 2000). These new mining concessions also overlap strongly with International Bird and Biodiversity Areas, another strong indicator of biodiversity (Fig. 1B).

**Figure 1.**
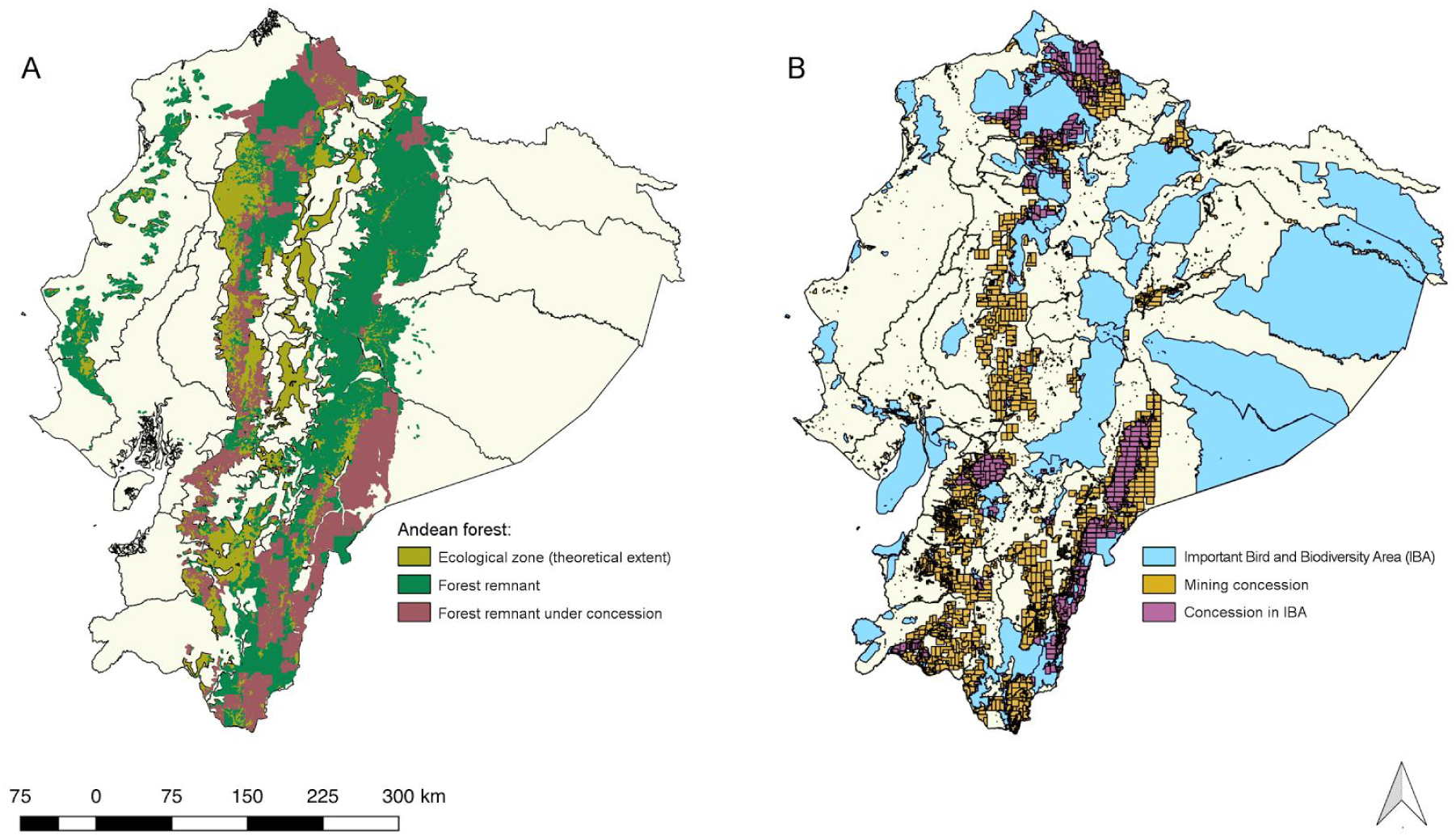
Maps showing the overlap between mining concessions and extant Andean forests and important bird and biodiversity areas (IBAs). In **A**, the Andean forest zone is shown, with deforested areas (yellow green), existing forest (dark green), and existing forest under mining concession (plum red). In **B**, mining concession are shown in yellow; Important Bird and Biodiversity Areas (IBAs) are shown in blue, and the overlap of these concessions with IBAs is shown in purple. Used with permission (Vandegrift, Thomas, Roy, & Levy, 2017).

Both mining and mining exploration decrease biodiversity through deforestation (Fig. 2A) and increased access and disturbance brought by the associated new roads (Asner et al., 2010; Bruijnzeel, 2004; Gross, 2017; Sonter et al., 2017). Forest cover is of key importance for both water quantity and quality because forests capture water, purify it, slow its movement through the landscape, and are themselves important for generating the clouds that produce the rain (Brauman, Daily, Duarte, & Mooney, 2007; Bruijnzeel, 2004; Foley et al., 2005). Water quality is best measured by aquatic macroinvertebrates because they live in the water and integrate both its physical and chemical environments (Rios-Touma, Acosta, & Prat, 2014). A recent study in Ecuador showed that water quality was excellent in Andean streams only when the headwater catchments had undisturbed native vegetation cover of >70% (Iniguez-Armijos, Leiva, Frede, Hampel, & Breuer, 2014).

**Figure 2.**
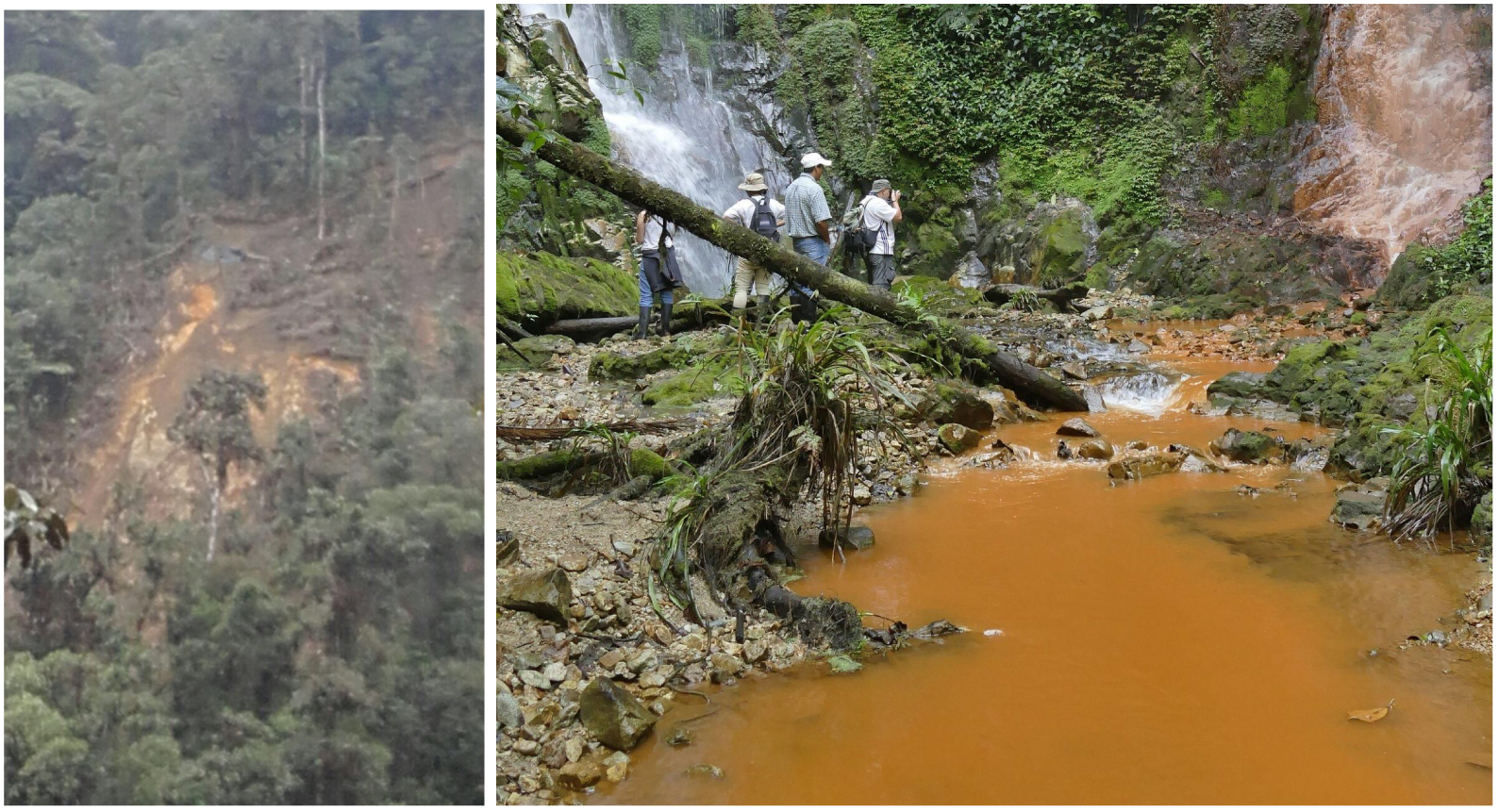
Consequences of mining exploration. **A**. Image of deforestation and landslide associated with mineral exploration in Imbabura Province, Ecuador, taken in September 2017. Photographer: anonymous. **B.** Water quality degradation (waterfall to right compared to left) caused by CODELCO exploration activities in the Junín Community Cloud Forest Reserve in late 2017. Photographer: C. Zorilla.

Deforestation reduces inputs of leaf litter into streams, changing energy inputs to the streams shifting trophic structure toward algal-based autotroph systems in Montane Choco-Andean Streams (Encalada, Calles, Ferreira, Canhoto, & Graca, 2010), leading to diminished aquatic macroinvertebrate and fish assemblages (Allard, Popee, Vigouroux, & Brosse, 2016; Teresa & Casatti, 2012). Moreover, after deforestation, mercury mobilization of soil is the main source of methylated mercury in aquatic systems in the northern Amazon basin (Roulet et al., 2000), with enormous negative effects on aquatic life and human health (Webb, 2005). Landslides, soil loss, increases in stream sediments, and changes in stream flows are additional problems that result from deforestation, see Figure 2 (Molina, Vanacker, Balthazar, Mora, & Govers, 2012; Restrepo, Kettner, & Syvitski, 2015; Roering, Schmidt, Stock, Dietrich, & Montgomery, 2003).

### Ecuador is the hottest hotspot of biodiversity in the world

The tropical Andes of Ecuador are at the top of the world list of biodiversity hotspots in terms of vertebrate species, endemic vertebrates, and endemic plants (Myers et al., 2000). The dominant pattern of biodiversity is to increase towards the equator and to decrease towards the poles (Brown, 2014). About half of all plant species occur in tropical forests near the equator, in only about 7% of the world’s total land surface area (Eiserhardt, Couvreur, & Baker, 2017). Tree diversity is highest in the tropical lowlands of the “Amazonas” part of South America, including Ecuador’s Oriente, whereas non-tree vascular plant diversity is concentrated in the highly dissected mountainous terrain and cloud forests of Northwestern South America, largely due to the high levels of endemism in such terrain (Gentry, 1992; Jørgensen & Léon-Yánez, 1999; Leon-Yanez et al., 2012; Ulloa et al., 2017).

Although biodiversity typically decreases with elevation, there is a secondary increase in diversity in the Andean cloud forest zone, which occurs between about 800 and 3500 m, depending on the distance from the sea and size of the mountain (Bruijnzeel, Mulligan, & Scatena, 2011). These persistently foggy and rainy forests are speciose with epiphytes, such as orchids and bromeliads (Gentry & Dodson, 1987; Kuper, Kreft, Nieder, Koster, & Barthlott, 2004). Several other taxa have higher diversity in the cloud forest zone relative to either higher or lower elevations, including: moths (Brehm et al., 2016), frogs (Willig & Presley, 2016), caddisflies (Blanca Ríos-Touma, Holzenthal, Huisman, Thomson, & Rázuri-Gonzales, 2017), and tree ferns (Ramirez-Barahona, Luna-Vega, & Tejero-Diez, 2011). Ecuador is crossed by two main mountain ranges, the Cordillera Occidental and the Cordillera Oriental, each with cloud forest zones that differ in floristic composition and that harbor specialized microhabitats with narrow endemic species and due to its latitude, Ecuador’s vegetation has northern and southern elements (Jørgensen & Léon-Yánez, 1999).

Tropical regions are diverse, to a large degree, because there is a constant supply of energy from the direct angle of the sun at the equator combined with high rainfall as a result of the heating (Kreft & Jetz, 2007). Water supply and energy drive about 70% of the variation in species diversity (Kreft & Jetz, 2007). However, some diversity can also be ascribed to habitat stability. Both fossil and phylogenetic methods suggest that some lineages have been present for 67–115 million years, indicating climate stability and low overall extinction rates (Eiserhardt et al., 2017). To the stability of climate at the low elevations, the Andes added rapid change in the uplands. Recent uplift, steep elevation and climate gradients over short distances, and spatial complexity, have led to vast changes in species composition on short spatial scales (Kreft & Jetz, 2007). The rapidity of the uplift of the Andes has also increased speciation rates by forming physical and climatic barriers to gene flow, and by opening up new niches (Antonelli, Nylander, Persson, & Sanmartin, 2009; Bell, 2004; Eiserhardt et al., 2017; Hughes & Eastwood, 2006; Kreft & Jetz, 2007; Scherson, Vidal, & Sanderson, 2008)Thus, the neotropics are acting both as a ***museum of biodiversity*** accumulated over a long time in the lowlands, and as a **c*radle of new innovations*** and speciation spurred by the uplift of the Andes (Kreft & Jetz, 2007).

### A Fragile Diversity

The spatial complexity that is partially responsible for Ecuador’s hyperdiversity also represents a particular vulnerability to land changes such as those posed by the proposed mining projects. Many Andean species have very limited ranges due to a combination of microclimatic and topographical barriers reducing dispersal (Eiserhardt et al., 2017; Hughes & Eastwood, 2006). For example, 27% of the known plants in Ecuador are endemic, and many of the endemics are known from only one of a few localities in a single province, and are thus not found anywhere else in the world (Jørgensen & Léon-Yánez, 1999; Leon-Yanez et al., 2012; Valencia, Pitman, León-Yánez, & Jorgensen, 2000). The rates of endemism are greater in the mountains than in the lowlands that straddle them (Borchsenius, 1997; Pitman & Jorgensen, 2002). With such spatially limited endemism, even a single mining project threatens the survival of species, such as the critically endangered longnose harlequin frog (*Atelopus longirostris*), which is in danger of extinction by the Llurimagua mining project (Tapia, Coloma, Pazmiño-Otamendi, & Peñafiel, 2017).

### Protected Lands in Ecuador

There are several major types of protected areas in Ecuador (Horstman, 2017; Lopez-Rodriguez & Rosado, 2017).

1. Heritage Natural Areas (= Patrimonio de Áreas Naturales del Estado, or Sistema Nacional de Área Protegidas = SNAP), including National Parks, which are set aside and funded by the Ecuadorian national government and run as public institutions.
2. Areas of Forest and Protected Vegetation (= Áreas de Bosque y Vegetación Protectora = ABVP = Bosques Protectores = BP), which are recognized by the national government but not funded by it. Recognition by the government of Bosques Protectores enables legal support when conflicts in land use occur, including help with illegal logging and squatters (Horstman, 2017).
3. Private Reserves, which are not necessarily recognized by the national government, nor funded by it. However, some national programs exist to promote the conservation of forests by private landowners, such as the successful Socio Bosque program. These are owned by individuals or private collectives.
4. Community Reserves, which are neither recognized by the national government, nor funded by it. These are “private” reserves that are owned and managed by local Parish governments for the benefit of the community.

There are numerous habitats and associated biodiversity that are underrepresented in the SNAP system, but three stand out in particular as needing more protection: coastal dry forests, which are located near population centers (Horstman, 2017), and the forests of southern and western Ecuador (Borchsenius, 1997; Sierra, Campos, & Chamberlin, 2002). The forests of the west, including cloud forests, are nearly gone. In 2000, it was estimated that more than 96% of the primary forested land in western Ecuador had been cleared (Myers et al., 2000), and much of that remaining 4% has been lost since then. A large portion of the remaining forest is in Bosques Protectores, and now 30% of these are under threat due to new mining concessions (Vandegrift et al., 2017). Figure 1A shows how the concessions disproportionately affect the southern and northwestern regions of the Andes, the areas with the highest biodiversity.

Bosques Protectores arose in the late 1980’s, with the enactment of the National Forestry Law (Horstman, 2017). While typically smaller than the nationally protected SNAP areas, Bosques Protectores are often relatively large (averaging 13,155 ha), and in total they currently make up about one third of protected lands in Ecuador (Vandegrift et al., 2017). Because they cover a wide diversity of habitats, even small Bosques Protectores are of great importance for protecting a diversity of endemic species, which are typically found at only a few localities (Borchsenius, 1997). In addition to Bosques Protectores, the Ministry of the Environment manages the Socio Bosque program, where landowners are paid up to $30 per hectare to conserve native forests on their land. Private reserves and community reserves often fail to qualify for formal status as protected areas, but represent a significant portion of conserved land in Ecuador. In the Intag valley, the local organization DECOIN (Organización de Defensa y Conservación Ecológica de Intag) has helped 38 communities purchase and manage community reserves, leading to the protection of some 12,000 hectares (28,650 acres) of land (Veintimilla 2017), including the Júnin Community Cloud Forest Reserve (discussed below).

Deforestation accounts for 12–24% of greenhouse gas emissions due to human activities (Gibbs, Brown, Niles, & Foley, 2007; IPCC, 2014). The Socio Bosque program pays communities or individuals to preserve forest to reduce climate change. For the last several years, Ecuador has been moving to increase the value of standing forests by investing in the Socio Bosque program, with the financial help of REDD+ (Lima, Visseren-Hamakers, Brana-Varela, & Gupta, 2017). More than 173,000 Ecuadorians have benefited from this program (Lima et al., 2017); however, recent ministry budgetary constraint has resulted in members failing to receive payments since 2015, with many questioning the program’s survival (Ortiz, 2017). This, in addition to the new mining concessions, indicate that the government is turning away from conservation.

### Mining and Environmental Legislation

Metal mining in Ecuador has historically been small-scale and artisanal, the majority of it concentrated in the south of the country. Ecuador’s mining legislation was correspondingly rudimentary and was not well defined until 1937, when subsoil metals were named property of the state. Environmental legislation specific to mining was absent from Ecuador until new laws came into effect in 1991 (Congreso 1991). This legislation limited the granting of concessions in protected lands and mandated environmental impact assessments for all mining activities. In 1994 the World Bank funded the Project for Mining Development and Environmental Control (Spanish acronym: PRODEMINCA) with the aim of developing the Ecuadorian mining sector (Davidov 2013). The project collected mineralogical information from 3.6 million hectares of mostly western Ecuador, including seven protected regions. The regulatory recommendations made by PRODEMINCA were codified into law in 2000, identifying mining as a national priority and significantly deregulated the sector (Congreso Nacional, 2000). However, under the revamped regulations, mining development remained prohibited in government protected areas (Tarras-Wahlberg et al., 2000), which have thus far been interpreted to be only the SNAP protected areas described above, leaving the Bosques Protectores vulnerable.

The next major changes occurred with the adoption of Ecuador’s new constitution in 2008, which included the “Mining Mandate” that reverted the majority of mining concessions to state ownership (Wacaster, 2010). The new constitution also included the historic decision to give nature inherent rights (articles 71–74, (Asamblea Nacional, 2008)). However, the new laws also allowed mining in protected areas by special request of the president and approval by the National Assembly. In 2009 the government of Rafael Correa authored a new Mining Law, which increased regulation on mining companies. While the law did augment some environmental standards, it was met by widespread protests by indigenous and social movements that had hoped for stronger environmental and social guarantees. In 2015 and 2016 the Correa government made deregulatory modifications to the mining law to incentivize foreign investment. These changes included decreasing the corporate tax rate and windfall tax on mining companies (Unda, 2017). This made the acquisition of mining concessions much easier, leading to the bidding and auctioning of mineral concessions in State possession throughout that year (Ministry of Mines, 2016), and resulting in the recent increase in granted concessions (Figure 1).

### A more sustainable way forward

Responsible development of the region’s infrastructure with an eye for long-term sustainability, education, ecotourism, and research represents a much more sustainable way forward for Ecuador’s last uncut forests, and the people who call them home (Asquith, Vargas, & Wunder, 2008; Kocian, Batker, & Harrison-Cox, 2011; Pozo, Aguirre, & Sanchez, 2016; Welford & Barilla, 2013). In fact, stable local businesses already exist adjacent to many Bosques Protectores and ecotourism in the area has experienced steady growth (Kocian et al., 2011). These local businesses promote ecotourism and science, and typically involve many community members of all ages and genders. This is in sharp contrast to the effects of mining, which typically creates a short-term economy that ends when the mines close, and with 95% of the jobs being held by men (Walter, Tomás, Munda, & Larrea, 2016).

In the rest of this review, we illustrate the major role that Bosques Protectores are playing in preservation of biodiversity and related ecosystem services, while also serving as a sustainable engine for local economies. To illustrate the biodiversity, we have built comparative species lists from several reserves in the exceptionally biodiverse Chocó and Tropical Andes regions of NW Ecuador. We also briefly indicate how each reserve is benefiting the local economy.

## METHODS

### Localities

The medium and large Bosques Protectores discussed herein are shown in Figure 3. They lie just to the South of the Cotacachi-Cayapas Ecological Reserve, and include: Los Cedros (68% in concession), El Chontal (95% in concession), Mashpi (96% in concession), and Maquipucuna (36% in concession). Since BP El Chontal is so poorly explored (virtually no published data), we used several other small reserves (<500 hectares) in the Intag Valley from which we could find data to gain an idea of the biodiversity in that region (BP La Florida Cloud Forest Reserve (abbreviated La Florida), El Refugio de Intag Lodge (abbreviated El Refugio), and the Júnin Community Cloud Forest Reserve (abbreviated Junín), see Figure 4. Hereafter, we will refer to this set of reserves as the “Intag”.

**Figure 3.**
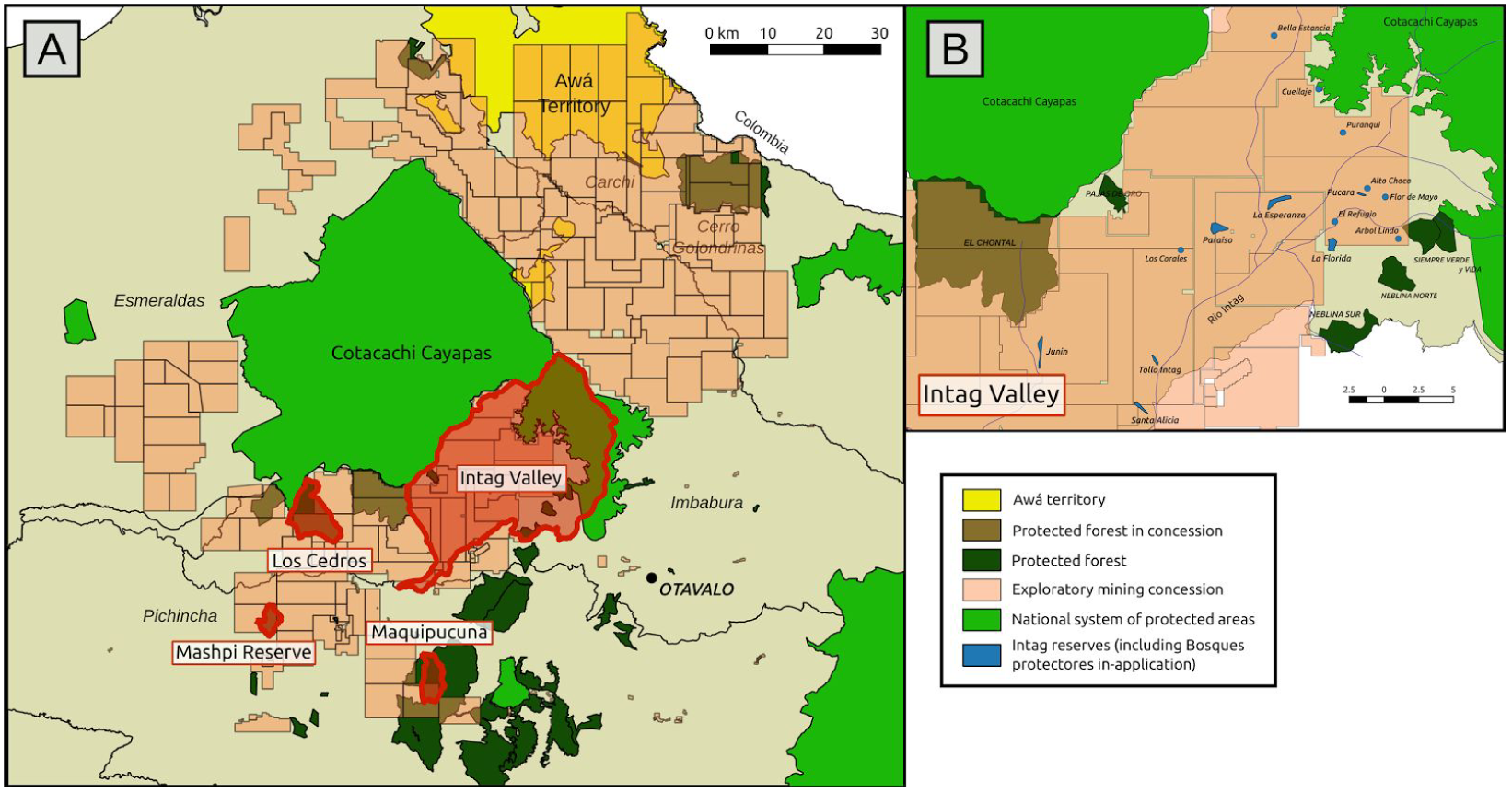
Maps showing the extent of the mining concessions and overlap with Bosques Protectores and Indigenous lands in the region around the Cotacachi-Cayapas Ecological Reserve. A. The Indigenous Awá lands (yellow) are to the north and are covered by almost 70% concessions, indicated in darker yellow; indigenous lands are discussed in a different publication (Vandegrift et al. 2017). All the Bosques Protectores under discussion herein are to the S of Cotacachi; they are dark green, unless covered by a concession, then they are brown. B. An expanded panel of the Intag Valley showing the smaller reserves in blue.

All the reserves studied herein lie in the region that is recognized to be the most important for the conservation of two critically endangered species, the brown-headed spider monkey (*Ateles fusciceps* ssp. *fusciceps*) (Peck et al., 2010), and the black-breasted puffleg hummingbird (*Eriocnemis nigrivestis*) (Jahn, 2008). It is also home to hundreds of other endangered species from birds to frogs to orchids (Tables 1–4 and Appendices 1–6), and includes Important Bird and Biodiversity Areas (Figure 1B). Two of the reserves, Los Cedros and El Chontal, share a border with the nationally protected Cotacachi-Cayapas Ecological Reserve.

**Table 1.** The 163 rare species known to occur at Bosque Protector Reserva Los Cedros as of 15 Jan 2018. Orange color indicates the rare classes, in order of most endangered, as defined by the International Union for Conservation of Nature (IUCN). Unique species are those not found at any of the other areas we studied.

| GROUP | (CR) | (EN) | (VU) | (NT) | (LC) | (DD) | (NE) | total | unique |
| --- | --- | --- | --- | --- | --- | --- | --- | --- | --- |
| Orchids<sup>a</sup> | 0 | 2 | 57 | 11 | 16 | 2 | 95 | 184 | 106 |
| Birds<sup>b</sup> | 0 | 3 | 9 | 13 | 276 | 0 | 1 | 302 | 22 |
| Mammals-Non-bat | 2 | 2 | 9 | 4 | 13 | 2 | 0 | 32 | 7 |
| Mammals-Bats<sup>c</sup> | 0 | 0 | 0 | 0 | 1 | 0 | 0 | 1 | 0 |
| Reptiles<sup>c</sup> | 0 | 0 | 1 | 4 | 4 | 0 | 0 | 9 | 4 |
| Amphibians | 0 | 4 | 2 | 4 | 2 | 2 | 1 | 15 | 4 |
| Other plants<sup>d</sup> | 0 | 9 | 16 | 10 | 5 | 0 | 190 | 236 | - |
| TOTAL | 2 | 20 | 94 | 47 | 323 | 6 | 290 | 782 |  |

<sup>a</sup> Understudied; 400 orchid species likely occur there says expert C. Dodson.
<sup>b</sup> About 100 more are likely; to protect the reserve there are few trails and it is infrequently visited due to distance from Quito.
<sup>c</sup> Not yet studied at the reserve.
<sup>d</sup> Understudied; the expected number of plants is well over 2,000. The reserve has never been catalogued. The plants listed here were mentioned in studies for other things, such as monkeys, or were genera specifically targeted for study by a specialist.

**Table 2.**
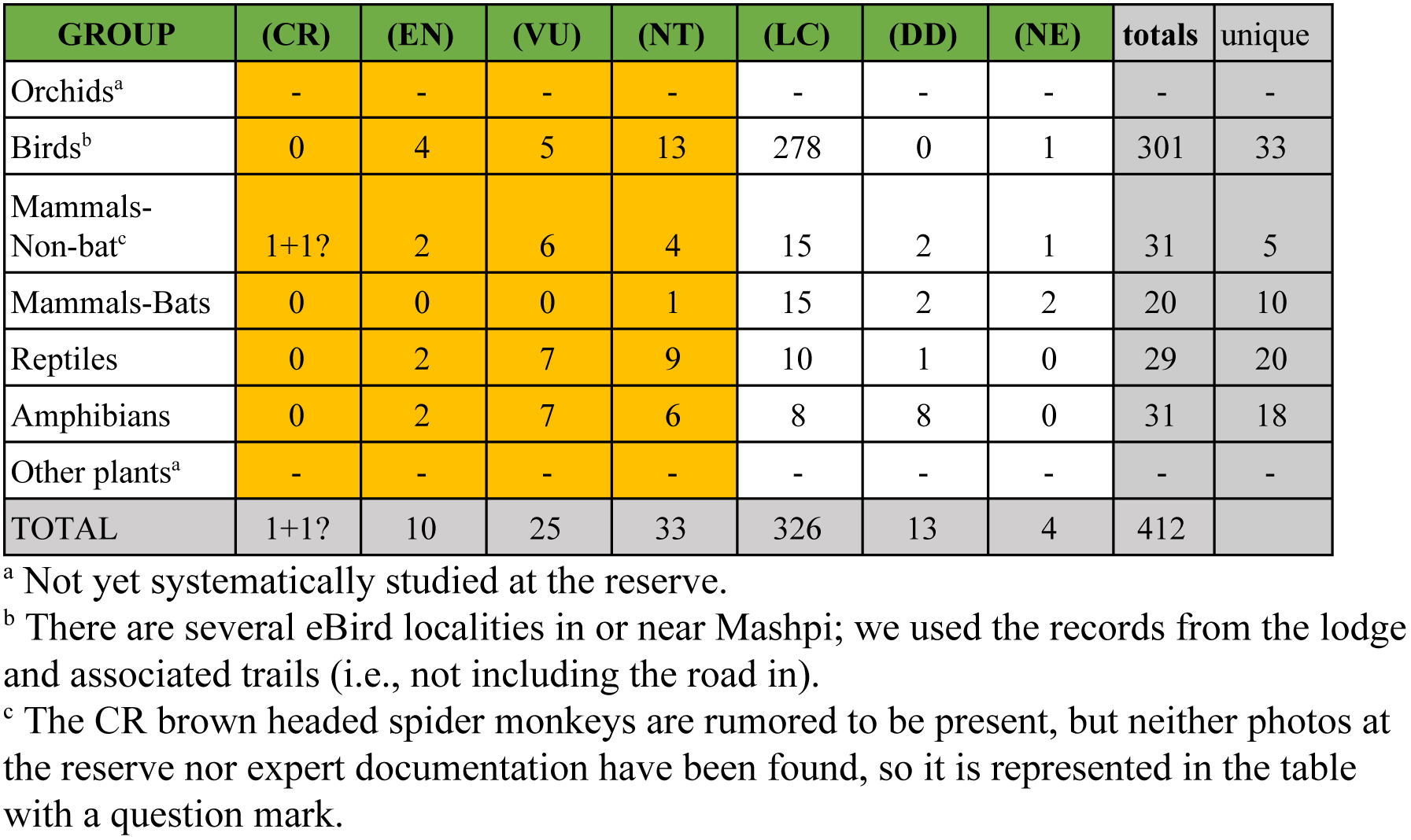
The 69 Rare species known to occur at Bosque Protector Mashpi as of 15 Jan 2018. Orange color indicates the rare classes, in order of most endangered, as defined by the International Union for Conservation of Nature (IUCN). Unique species are those not found at any of the other areas we studied.

**Table 3.** The 99 Rare species known to occur at Bosque Protector Maquipucuna as of 15 Jan 2018. Orange color indicates the rare classes, in order of most endangered, as defined by the International Union for Conservation of Nature (IUCN). Unique species are those not found at any of the other areas we studied.

| GROUP | (CR) | (EN) | (VU) | (NT) | (LC) | (DD) | (NE) | totals | unique |
| --- | --- | --- | --- | --- | --- | --- | --- | --- | --- |
| Orchids | 0 | 1 | 45 | 14 | 16 | 2 | 204 | 282 | 207 |
| Birds | 0 | 1 | 4 | 6 | 296 | 0 | 0 | 307 | 18 |
| Mammals-Non-bat | 0 | 2 | 4 | 5 | 15 | 4 | 0 | 30 | 9 |
| Mammals-Bats | 0 | 0 | 0 | 0 | 11 | 0 | 0 | 11 | 3 |
| Reptiles | 1 | 0 | 2 | 4 | 6 | 2 | 2 | 17 | 6 |
| Amphibians | 0 | 5 | 3 | 2 | 4 | 1 | 2 | 17 | 5 |
| Other plants<sup>a</sup> | - | - | - | - | - | - | - | 1714 | - |
| TOTAL | 1 | 9 | 58 | 31 | 348 | 9 | 208 | 2378 |  |

<sup>a</sup> A flora has been completed (Webster & Rhode, 2001, 2005), but with the exception of the orchids, we did not individually query each of the 1,996 species in Tropicos for rarity; quite a few will be rare.

**Table 4.**
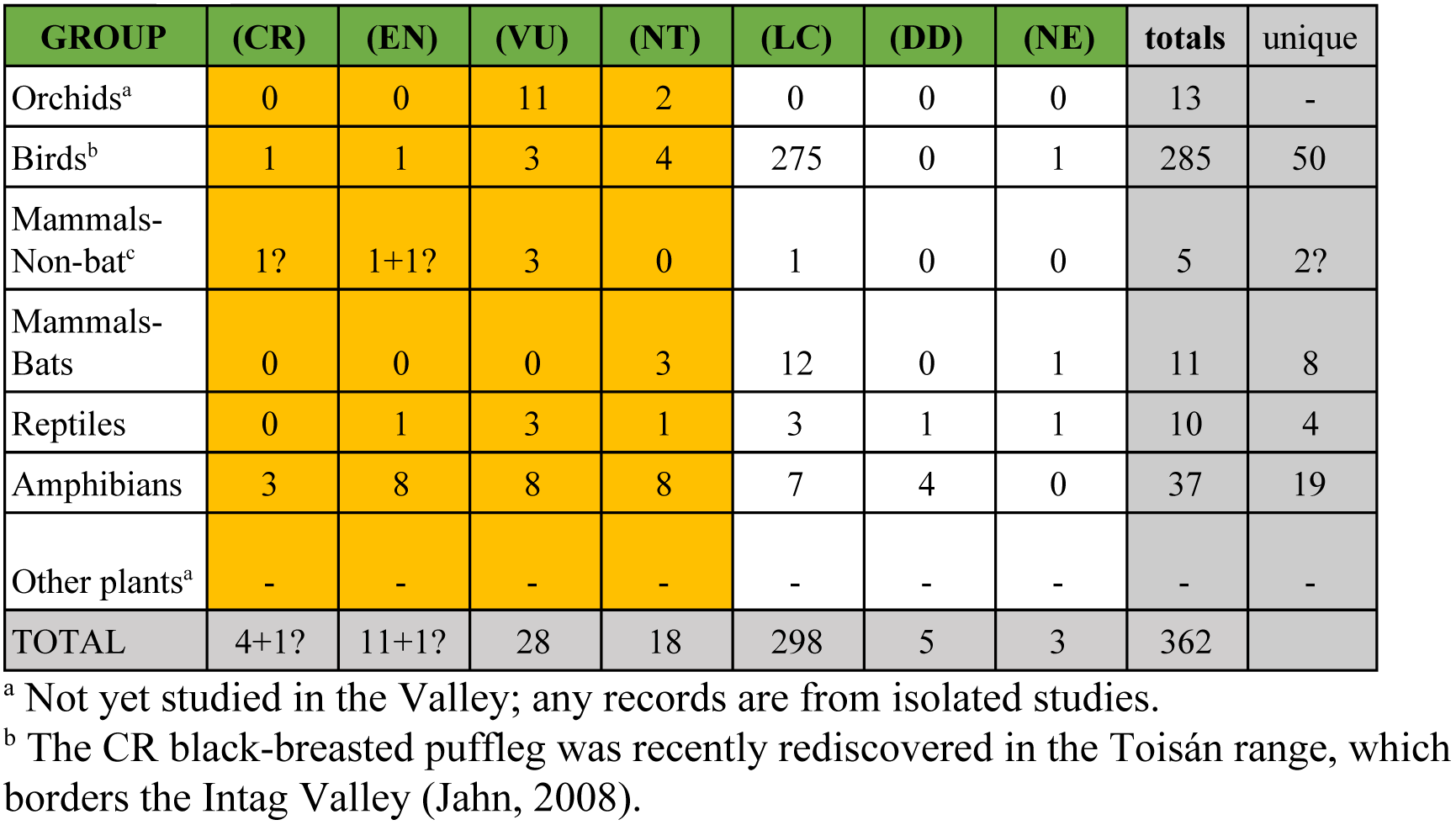
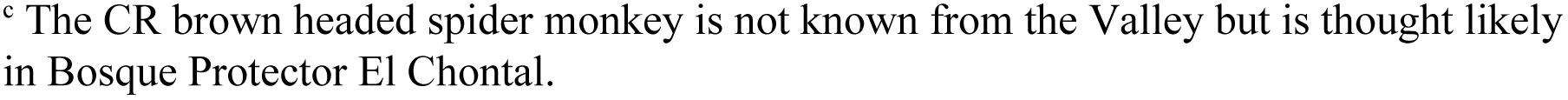
The 61 Rare species known to occur in the Intag Valley as of 15 Jan 2018. Orange color indicates the rare classes, in order of most endangered, as defined by the International Union for Conservation of Nature (IUCN). Unique species are those not found at any of the other areas we studied.

| GROUP | (CR) | (EN) | (VU) | (NT) | (LC) | (DD) | (NE) | totals | unique |
| --- | --- | --- | --- | --- | --- | --- | --- | --- | --- |
| Orchids<sup>a</sup> | 0 | 0 | 11 | 2 | 0 | 0 | 0 | 13 | - |
| Birds<sup>b</sup> | 1 | 1 | 3 | 4 | 275 | 0 | 1 | 285 | 50 |
| Mammals-Non-bat<sup>c</sup> | 1? | 1+1? | 3 | 0 | 1 | 0 | 0 | 5 | 2? |
| Mammals-Bats | 0 | 0 | 0 | 3 | 12 | 0 | 1 | 11 | 8 |
| Reptiles | 0 | 1 | 3 | 1 | 3 | 1 | 1 | 10 | 4 |
| Amphibians | 3 | 8 | 8 | 8 | 7 | 4 | 0 | 37 | 19 |
| Other plants<sup>a</sup> | - | - | - | - | - | - | - | - | - |
| TOTAL | 4+1? | 11+1? | 28 | 18 | 298 | 5 | 3 | 362 |  |

<sup>a</sup> Not yet studied in the Valley; any records are from isolated studies.
<sup>b</sup> The CR black-breasted puffleg was recently rediscovered in the Toisán range, which borders the Intag Valley (Jahn, 2008).
<sup>c</sup> The CR brown headed spider monkey is not known from the Valley but is thought likely in Bosque Protector El Chontal.

### Data Collection

Species lists were assembled for all the reserves for mammals, birds, amphibians, and reptiles (Appendices 1–4). The orchids were assembled for the two reserves with specimen vouchered data (Maquipucuna and Los Cedros, Appendix 5), and a partial list of plants was assembled for Los Cedros, primarily from published papers, but also including some previously unpublished data (Appendix 6). Nomenclature follows that of the Tropicos (2017) plant database.

To assemble the bird table, we used records from eBird for each of the localities for which these lists existed (Los Cedros (eBird, 2017c), La Florida (eBird, 2017b), Maquipucuna (eBird, 2017d), Mashpi (eBird, 2017e), and El Refugio (eBird, 2017a). eBird is vetted by local experts who verify the occurrences, and it uses a standardized format. We added in any published bird data found, if it was not yet in eBird, and additional data from the reserve managers at La Florida and El Refugio. For bird common names in English, we used eBird, for common names in Spanish we used the *Lista de las Aves del Ecuador* (J. F. Freile et al., 2015-2017). To assemble the other species lists we used reserve records when backed up by photos, videos, or experts, and to search for publications we used the reserve names as keywords in Google Scholar, Web of Science, and the Google search engine. We also searched for protected area place names in the excellent online databases for amphibians (Ron, Yanez-Muñoz, Merino-Viteri, & Ortiz, 2017), reptiles (Torres-Carvajal, Pazmiño-Otamendi, & Salazar-Valenzuela, 2017), and mammals (Brito, Camacho, Romero, & Vallejo, 2017) produced by the Museo de Zoología, Pontificia Universidad Católica del Ecuador (PUCE). We used the Spanish common names for all animals from these databases, and the English common names from either the Museo de Zoología websites or IUCN (2017).

### To assess rarity

We used a combination of international and national (Ecuadorian) databases. For birds and animals, we used the *IUCN Red List of Threatened Species* (IUCN, 2017), and for the plants, the Tropicos database (Tropicos, 2017). Since the international databases are not updated as regularly as the Ecuadorian Red lists as cited in the Museo de Zoología databases (Brito et al., 2017; Ron et al., 2017; Torres-Carvajal et al., 2017), we give both assessments when they differ, with the national red list proceeding that of the IUCN in our lists. Most of the rare species in Ecuador are endemics that do not occur anywhere else, so the Ecuador Red list is usually the most accurate rarity assessment. All the databases use the IUCN graduated system of rarity (IUCN, 2017), ranging from least threatened to extinct: NT=near threatened, VU=vulnerable to extinction, EN=Endangered, CR=Critically endangered, EW=Extinct in the wild, EX=Extinct. There are three other categories not included as rare in our lists: LC=least concern, DD=data deficient, and NE=never evaluated. We note, however, that quite often organisms that are in the DD or NE categories are also rare, they simply have not been assessed yet (NE) or there is insufficient data to assess them (DD), which is often an indication of rarity.

## RESULTS

### Los Cedros (0°18’35.62”N, 78°46’47.01”W)

(http://reservaloscedros.org) is located between 980 and 2,200 m elevation. It is fully in the cloud forest zone. It receives 2903±186 mm of rain per year at the 1,300 m elevation of the fieldstation, based on 15 years of reserve records (J. DeCoux, pers. comm.). At the higher elevations, considerably more rain falls. Los Cedros is remote; it takes 6–7 hours to get to Los Cedros from Quito, including a two-hour mule ride. Sixty-eight percent of its 5,256 hectares of protected cloud forest have recently been put into mining concessions. Candidate areas for copper-containing porphyries in the reserve have been identified by aeromagnetic surveys, conducted without permission of the landowners. Los Cedros is not accessible by road, and for this reason has been, to date, both better protected and less scientifically explored than some other Bosques Protectores.

#### Species

Los Cedros is known to protect at least 163 rare species (critically endangered = 2, endangered = 20, vulnerable = 94, and near threatened = 47) see Table 1 and Appendices 1–6). Its remoteness is why it still has three species of monkey: the critically endangered brown-headed spider monkey (*Ateles fusciceps fusciceps),* the vulnerable white-headed capuchin (*Cebus capucinus*), and the endangered mantled howler monkey (*Alouatta palliata*), as well as the vulnerable Andean Spectacled Bear (*Tremarctos ornatus*) (Appendix 1). Remoteness and good habitat also explain why there are six species of cats, including the critically endangered Jaguar (*Panthera onca*), the vulnerable Oncilla (*Leopardus tigrinus*) and the near threatened Margay (L*eopardus wiedii*). Jaguars are now extremely rare in western Ecuador due to habitat loss and need for large ranges (Zapata-Ríos & Araguillin, 2013)In addition to Los Cedros (BirdLife International, 2017), a Jaguar was recently photographed in nearby (<5 km) Manduriacu Reserve (Jost, 2016)on the Manduriacu river, which originates in Los Cedros, and have also been seen in the adjacent Cotacachi-Cayapas Ecological Reserve (Zapata-Ríos & Araguillin, 2013). Prey include the Little Red Brocket Deer (vulnerable), which—along with other prey such as the agouti, peccary, and monkeys—are rapidly hunted out of reserves by people when there are nearby roads.

Los Cedros is a bird hotspot (eBird, 2017c). Of the 302 bird species seen at Los Cedros (Appendix 2), at least 25 are endangered, vulnerable, or near threatened due to habitat loss, even before the latest mining concessions. Many of the birds at Los Cedros are found only in the cloud forests of the Chocó region (BirdLife International, 2017; Cooper, Ridgely, Ortiz, & Jahn, 2006), and include very recently described species such as the cloud-forest pygmy owl (*Glaucidium nubicola*) (J. Freile et al., 2013). In addition, these forests harbor a number of vulnerable and near threatened neotropical migrants that summer in Canada and the United States, such as the cerulean warbler (*Setophaga cerulea)* and olive-sided flycatcher (*Contopus cooperi)*, whose populations depend on having winter habitat. Comparing the reserves being highlighted here, twenty-two species of bird are only found at Los Cedros and not at the other reserves, including 5 of the 25 rare birds (Appendix 2). Based on the number of reported species in nearby reserves, and habitats present, it is expected that the final list for Los Cedros will have around 400 bird species; it is less frequently “birded” than more accessible reserves.

The frogs are fantastic, almost all rare and found only in the local cloud forests (Appendix 3). For example, the recently described rainfrog, *Prisimantis mutabilis,* is only known from two streams, one of which is at Los Cedros (Guayasamin, Krynak, Krynak, Culebras, & Hutter, 2015). This remarkable frog is able to change its skin texture, a feature never before seen in frogs (Guayasamin, Krynak, et al., 2015). Another of the rainfrogs was described from and named for Los Cedros: *Pristimantis cedros*. This species is locally common at Los Cedros, but has not been collected elsewhere (Hutter & Guayasamin, 2015).

Reptiles (Appendix 4) and bats (Appendix 1) are yet to be systematically studied at Los Cedros, but incidental records indicate that the reptiles are likely to be interesting. For example, there are coral snakes (*Micrurus ancoralis*, NT) and their mimics (*Oxyhopus petolaris*, LC) and the bizarre reticulate worm snake (*Amerotyphlops reticulatus*), which looks like a very fat foot-long worm and lives in the litter layer of the forest. Bats tend to be widespread without local endemics as can be seen in Table 2, but it would still be useful to determine which bats are at Los Cedros.

The Los Cedros forest is extraordinarily rich in plant species, with at least 299 tree species per hectare (Peck et al., 2010; Thomas, Vandegrift, Ludden, Carroll, & Roy, 2016). Associated with this forest are many fungi (Dentinger & Roy, 2010; Policha et al., 2016; Thomas et al., 2016), which are essential for forest growth (Vandenkoornhuyse, Quaiser, Duhamel, Le Van, & Dufresne, 2015) and decomposition (Yang et al., 2016). Two species of fungi proposed to the relatively new IUCN Global Fungal Red List Initiative, *Lamelloporus americanus* and *Hygrocybe aphylla*, are known from Los Cedros (Newman, Vandegrift, Roy and Dentinger, unpublished data), and many additional rare taxa are anticipated. Collections made there since 2008 have resulted in several hundred morphospecies, whose precise identifications are the subject of ongoing research.

Many plants in the Los Cedros forest are local endemics with small ranges (Appendix 6), including several orchids only known from Los Cedros (Appendix 5). Los Cedros currently has 187 orchid species on its list (Appendix 5). Of these, 71 (38.5%) are known to be some category of rare (endangered, vulnerable, threatened) and most of these are localized endemics. Seventeen of these rare orchids were originally described from Los Cedros, and at least seven of these have never been found elsewhere. Ninety-eight (52%) of the orchid species from Los Cedros have never been evaluated for rarity because they are not endemics and it is difficult to assess rarity across country borders (Endara & Jost, 2011). However, we note that at least a dozen of the NE species barely range into Colombia, and are thus likely threatened.

The numbers of orchid species found to date at Los Cedros (Appendix 5) are underestimates because of its inaccessibility; the final list is likely to be near 400 species (C. Dodson pers. com.). The absolute size of the orchid floras cannot be compared with our data, since the orchids have not been completely catalogued at Los Cedros, but we could examine the overlap of what was known at Los Cedros with the only other reserve for which orchid data was available, the better studied Maquipucuna. Los Cedros shares only 43% of its known orchid diversity with Maquipucuna. For a specific example, there are 14 species in the orchid genus *Dracula* at Los Cedros (Appendix 5), all of but three of which are endangered or vulnerable. Only four of the 14 *Dracula* species at Los Cedros also occur at Maquipucuna, at which only five species of *Dracula* have been recorded (Appendix 5 and (Webster & Rhode, 2001)).

Note that each orchid species is associated with pollinators, which themselves are speciose and understudied. For example, studies of the mushroom-mimicking orchid *Dracula lafleurii* uncovered at least 60 new species of fruit flies that pollinate it (Endara, Grimaldi, & Roy, 2010; Policha, 2014; Policha et al., 2016; Policha et al., submitted). These unnamed flies are related to a model organism, the common fruit fly (*Drosophila melanogaster*), which is widely used in genetics and neurobiology studies that benefit humans (Roberts, 2006).

Los Cedros protects the origins of three rivers: the Río Manduriacu, the Río Verde, and the Río Los Cedros, plus it encompasses the south bank of the upper Río Magdalena Chico. These rivers supply freshwater to people lower down, and are the habitat for an amazing diversity of life themselves. In an exploratory three‐ night survey, almost 40 species of caddisflies (Trichoptera) were collected, of which more than a third are probably new to science (Blanca Ríos-Touma et al., 2017). Considering that this is only one of the eleven orders of aquatic macroinvertebrates in the area (Knee & Encalada, 2014), the potential number of novel species is enormous.

#### Social & Economic

The field station at Los Cedros can lodge up to 40 people at a time, and is visited by local and university classes, ecotourists, and scientists. It benefits the nearby communities of Magdalena Alta and Chontal with employment (guides, cooks, etc.) and by buying supplies and services there. Ecuadorian visitors and scientists are charged lower rates than tourists and foreign scientists. Los Cedros is governed by Fundación Los Cedros, which includes staff, local community leaders, and representatives from environmental groups.

### Mashpi (0° 9’57.17”N, 78°52’39.38”W)

(https://www.mashpilodge.com) is located between 550 and 1400 m elevation, and thus encompasses both tropical forest (to about 900 m) and lower montane cloud forest above that. The reserve website states Mashpi receives up to 6 meters of rain, but to our knowledge, there is not a weather station at the lodge. Mashpi is less remote than Los Cedros--about a three-hour drive from Quito. Ninety-six percent of its 1,178 hectares are now in mining concessions.

#### Species

Mashpi is known to protect at least 69 rare species (critically endangered = 1+1?, endangered = 10, vulnerable = 25, and near threatened = 33, see Table 2 and Appendices 1–5). The forest at Mashpi is still in excellent condition, as indicated by the presence of two primate species (the critically endangered white-fronted capuchin (*Cebus aequatorialis*) and the endangered howler monkey (*Alouatta palliata)*, as well as several cats (Table 2). Historically, Mashpi was part of the range for the critically endangered brown-headed spider monkeys (Peck et al., 2010), and though there are rumors that spider monkeys have been seen at Mashpi, we found no photos or expert sightings to verify this; we represent this uncertainty with a question mark in Table 2 and Appendix 1. The lower elevation of Mashpi compared with all the others we discuss in this region enables the presence of species that occur in warmer, lower elevation forests, such as anteaters (*Tamandua mexicana*).

Mashpi is also a bird hotspot (eBird, 2017e), with 301 species recorded from the lodge and trail system (Appendix 2). It protects a different set of birds than Los Cedros, with 33 unique species (Appendix 2), in part reflecting its lower elevation than the other reserves and its combination of montane tropical and lower cloud forests. Of the 22 rare & endangered bird species at Mashpi (Appendix 2), seven are not found at any of the other reserves examined. Perhaps the most interesting of these is the endangered Chocó vireo (*Vireo masteri*), which is only known from a few localities in Colombia and one, Mashpi, in Ecuador (BirdLife International, 2018).

About half of Mashpi’s observed amphibians, 15/31, are endangered, vulnerable, or near threatened, and about a third are endemic to Ecuador (Table 2, Appendix 3). The amphibians very clearly indicate the lower elevations at Mashpi. Eighteen of its 31 amphibians have thus far only been found there (Appendix 3) and not at the other reserves we are profiling, and all 18 of these have ranges mostly under 900 m (Appendix 3), the lower elevation limit for the other reserves included herein. Most of the lower elevation amphibians are widespread lowland forest “chocoan” species, whereas endemism and rarity is concentrated in the higher elevation cloud forest taxa (Appendix 3). Mashpi is the primary home for the Mashpi stream tree frog (*Hyloscirtus mashpi*), which was described from its streams. This frog is only known from a total of three localities and is most common at Mashpi (Guayasamin, Rivera-Correa, et al., 2015).

Of the reserves reported on here, Mashpi is the only one that has had dedicated attention paid to the reptiles, and thus its list is more complete: 29 species to date. Similar to the amphibians, many of the reptiles at Mashpi are reported from there and not the other reserves (20/29 or 69%, Table 2, Appendix 4). Warmer, lower elevations likely led to a higher number of species present, including the South American snapping turtle (*Chelydra acutirostris)* and the Northern eyelash boa (*Trachyboa boulengeri*), which do not occur at higher elevations. More than half the reptiles are rare, including two vipers, which are usually killed when humans encounter them.

Mashpi has also paid attention to its aquatic biodiversity. Preliminary results indicate there are at least 21 fish species (Franco, Falconí, Ríos-Tourma, Morochz, & Tobes, 2017) and up to 96 genera of aquatic macroinvertebrates (B. Ríos-Touma, Morabowen, Tobes, & Morochz, 2017), including around 60 species of caddisflies (Blanca Ríos-Touma et al., 2017).

#### Social & Economic

The ecolodge at Mashpi is a five-star hotel that has garnered international praise for its innovation and sustainability (Mashpi, 2018). As part of their commitment to sustainability, they use some of their profits to maintain a scientist on staff. Support of the local communities includes education opportunities, the hiring of guides and staff for the lodge and buying of supplies from local producers. Also, at San José de Mashpi, preserves like Mashpishungo and Pambiliño work in conservation, grow sustainable produce, and community empowerment through ecotourism.

### Maquipucuna (0° 7’0.12”N, 78°37’45.23”W)

(https://www.maquipucuna.org) is located between 900 and 2,700 m elevation. Thirty-six percent of its 2,474 hectares of protected land are now in mining concessions. Rainfall has never been systematically measured at Maquipucuna, but is likely to be at or above that of nearby Nanegalito (3230 mm) according to Webster and Rhode (2001). Of the reserves detailed here, Maquipucuna is the least remote; taking only two hours on developed roads from Quito. For this reason, it has more visitors and is better understood scientifically, but its wildlife and birds are adversely affected by the proximity to roads. For example, there are no longer monkeys at Maquipucuna.

#### Species

Maquipucuna is known to protect at least 99 rare species (critically endangered = 1, endangered = 9, vulnerable = 58, and near threatened = 31, see Tables 3 and Appendices 1–5). The most interesting mammal (Appendix 1) is the endangered Spectacled Bear (*Tremarctos ornatus*), the only South American bear, which also occurs at two of our other highlighted reserves, Los Cedros and in the Intag Valley. When the wild avocados are fruiting, the bears migrate to a few places where they are easily seen in Maquipucuna, creating a tourist attraction (Maquipucuna, 2018). About a third (9/30) non-bat mammals at Maquipucuna do not appear on the lists of any of our other studied reserves (Appendix 1). Of these, six are common LC species, but two are interesting near threatened small mammals, the water opossum *Chironectes minimus* and the mountain paca, *Cuniculus taczanowskii,* and one, the beady-eyed mouse, *Thomasomys baeops*, is data deficient.

Maquipucuna is also a bird hotspot (eBird, 2017d), with 307 species recorded from the lodge and trail system (Appendix 2). It protects a different set of birds than the other reserves, with 18 unique species. However, of the 11 rare and endangered bird species at Maquipucuna (Appendix 2), only one is unique to Maquipucuna, the black and chestnut eagle (*Spizaetus isidori)*. Some of the rare birds missing from Maquipucuna, but present at the other reserves, are the ground-dwelling birds, such as the Baudo guan, *Penelope ortini,* which suffer when nearby roads facilitate illegal hunting.

Ten of the seventeen amphibians reported from Maquipucuna are some category of rare, including eight species of rainfrog, one toad, and one salamander (Appendix 3). Similar to Los Cedros and Mashpi, Maquipucuna has a frog species, *Hyloxalus maquipucuna*, that was discovered there and is known only from this locality, but in this case, it is member of the poison dart frog family (Dendrobatidae) instead of being a rainfrog (Strabomantidae). Four other frog species from Maquipucuna are also not found at our other reserves (Appendix 3), following the pattern of localized cloud forest endemics.

Seventeen species of reptiles have been reported from Maquipucuna (Appendix 4). The most endangered species is an endemic snake, *Tantilla insulamontana,* which is critically endangered. Very little is known about this snake, which has been rarely seen; the main threats are habitat destruction, fragmentation and contamination (Torres-Carvajal et al., 2017).

Orchids (Appendix 5) were largely discussed under Los Cedros, the only other BP under discussion here for which we have detailed and at least partially vouchered data for orchids. Maquipucuna has one endangered orchid species, *Masdevallia ventricularia*, which is also at Los Cedros, and it has 45 vulnerable and 14 near threatened orchids (Table 3 & Appendix 5). Maquipucuna only shared 75 species with Los Cedros. While Los Cedros is particularly rich in *Dracula* and other pleurothallids such as *Acronia*, Maquipucuna is richer in *Cyrtochilum*, *Elleanthus* and *Epidendrum* species. These differences may be real due to topographic or climatic differences (the Río Guayllabamba runs between the reserves and could be a barrier, for example), or they may be due to collection bias at Los Cedros. We hope that our lists spur future work.

#### Social & Economic

Maquipucuna has an ecolodge frequented by birders and other ecotourists, and its website (Maquipucuna, 2018) states that “over 120 families benefit from ecotourism projects initiated and supported by Maquipucuna”. For example, they helped the nearby village of Yungilla to switch from charcoal production to reforestation and ecotourism (Gosdenovich, 2015; Houns, 2013), and are working to find ways to grow coffee and cacao more sustainably (Gosdenovich, 2015; Justicia, 2007).

### Intag Valley

The Intag Valley is a region in the Cotacachi canton of the Imbabura province, partially defined by its location as the watershed of the Intag river, but also defined culturally by the network of communities in eastern Cotacachi canton that cooperate on conservation and economic development projects. Due to earlier mining concessions and rich copper deposits, exploration has progressed the furthest in the Intag Valley as compared to elsewhere in NW Ecuador, with significant environmental consequences already apparent, just from “exploration” (Figure 2). We aggregated all data from the Intag area into a single column (“Intag”) in the Appendices, but kept the source of the data separate; most of what we found was from reserves 2–4, below, for location in the Intag Valley see (Fig. 3):

1. Bosque Protector El Chontal (0° 21’ 45’’ N, 78° 42’ 4” W) (http://www.zoobreviven.org/elchontal.htm) with an elevation range between 1,000 and 4200 m. Ninety five percent of its 6,989 hectares are now in mining concessions (Agencia de Regulación y Control Minero, 2017; Vandegrift et al., 2017).
2. Bosque Protector La Florida Cloud Forest Reserve (0° 22’ 0.01” N, 78° 28’ 54.17” W) (https://intagcloudforest.com) with an elevation range between 1800 and 2800 m. El Refugio de Intag Lodge (0° 22’ 25.32’’ N, 78° 28’ 33.6’’ W), (www.elrefugiocloudforest.com), a private reserve.
3. Júnin Community Cloud Forest Reserve (0° 17’ 18.11” N, 78° 40’ 0.47” W), a community owned reserve (http://www.junincloudforest.com).

#### Species

The Intag Valley is known to protect at least 61 rare species (critically endangered = 4+1?, endangered = 11+1?, vulnerable = 28, and near threatened = 18, see Table 4, & Appendices 1–5). The Intag Valley is home to four critically endangered species (Table 1): the black-breasted puffleg (*Eriocnemis nigrivestis)*, two frogs (*Ectopoglossus confusus* and *Hyloacalus jacobspetersi*), and a toad, confusingly called the harlequin frog (*Atelopus longirostis*). A fifth critically endangered species, the brown-headed spider monkey, is likely to be in the under-explored Bosque Protector El Chontal (Peck et al., 2010).

The only non-bat mammals that have been reported from Intag are all large, and all but one is rare (Table 4, Appendix 1), including the endangered Spectacled Bear (*Tremarctos ornatus*). Another endangered mammal that may be in El Chontal/Intag Valley, is the mountain tapir (*Tapirus pinchaque*). According to the El Chontal website (Fundación Zoobreviven, 2018) mountain tapirs are present and being hunted, but there are no photos and we could find no other modern records of this species being present in the Intag, so we represent the potential presence of this species with a question mark in Appendix 1. There is good data on the bat fauna of the Intag Valley, because they were mist netted in the Junín Cloud Forest Reserve (Cueva-A, Pozo-R, & Peck, 2013). The only other reserve that has comparable bat data is Maquipucuna (Appendix 1). None of the bats in Maquipucuna are rare, but three are near threatened (NT) in the Intag.

The Intag combined list of birds is 277, just a few short of the 300 eBird uses to define a “hotspot”. This list (Appendix 2) is likely incomplete since neither the extensive El Chontal Reserve nor the nearby Junín have been surveyed. The Intag provides homes for a different subset of birds than Los Cedros, Mashpi, or Maquipucuna, with the most unique species (50) (Appendix 2). Some of the difference in species from the other reserves may be due to the region’s proximity to drier, inter-Andean valleys, such as the beautiful jay (*Cyanolyca pulchra),* and others, such as the gray-breasted mountain toucan (*Andigena hypoglauca)* and the critically endangered hummingbird (*Eriocnemis nigrivestis*) reflect the high elevations (> 1500 m and up to 4,000 m) in this valley.

The amphibian fauna from the Intag is breathtaking, with an astonishing 26 species, mostly frogs, in some form of endangerment (Appendix 3), including three that are critically endangered. The majority of the rare amphibians in Ecuador are in the montane cloud forests, such as in the Intag, where the localized responses to small climate differences led to numerous speciation events and the formation of localized endemics (Arteaga et al., 2016). In 2016, researchers rediscovered the longnose harlequin frog (*Atelopus longirostris*) within the Junín Cloud Forest Reserve. This is an endemic species last seen in 1989 and previously listed as Extinct by the IUCN (Tapia et al., 2017). The rediscovery of the harlequin frog highlights the need for further research on amphibian diversity in the Intag.

Reptiles (Appendix 4) and orchids (Table 4) have not been well-studied in the Intag region, although the steep altitudinal gradients suggest the community of orchid species in Intag’s upper montane cloud forests are likely to differ significantly from those found in Los Cedros and other lower elevation forests (Gentry, 1992).

#### Social & Economic

El Chontal is run by the community Chalguayacu Alto, the Association Ganaderos y Agricultores, and Fundación Zoobreviven. The Junín Cloud Forest Reserve is owned and managed by a community organization that also manages a tourism business, the Ecocabañas Junín. Founded in 2000 with the help of DECOIN, the Ecocabañas provide an additional source of income for 40 local community members (Murillo & Sacher, 2017). The La Florida Cloud Forest Reserve is privately managed, and also supports the livelihoods of surrounding families via a tourism and education center. In addition to guiding and homestays the center provides environmental education to local and visiting students. Similarly, the El Refugío Lodge is a social enterprise that employs only local community members and supports cultural events in the town of Santa Rosa. The reserve managers and associated tourism operators regularly coordinate with local schools to host field trips, encouraging students to learn about their local watersheds as well as the wildlife that can be found within them.

## DISCUSSION

The reserves highlighted in this paper collectively protect an astonishing 269 rare species, including eight critically endangered species—of which two are primates (brown-headed spider monkey and white-fronted capuchin)—37 endangered, 140 vulnerable, and 84 near threatened species, as well as a very large number of more common species. Importantly, each reserve protects a unique subset of species that are not found at the other reserves. The reserves also serve their surrounding communities by providing sustainable jobs, which have gradually been increasing over time (Walter et al., 2016), and through ecosystem services such as clean and abundant water.

The still federally protected SNAP areas in Ecuador do not do a good job of protecting the localized endemics, which are scattered around the country (Endara, Williams, & León-Yánez, 2009). Furthermore, the SNAP system primarily protect the lower elevations of the eastern slopes of the Andes, missing the highly diverse mountains (Endara et al., 2009). We show here that a large number of endemics are currently being protected in Bosques Protectores, but that these are now endangered by mining. The BPs highlighted are near or adjacent to the Cotacachi-Cayapas Ecological Reserve, and are acting both as buffers and corridors for it, and they extend the protected elevation gradient. Cotacachi is largely at high elevation (>2,500 m) at its southern end. The BPs discussed here extend protection into the critically endangered NW cloud forest zone and upper montane forests that occur between 900-2500 m. These are the habitats preferred by the most endangered species in our study, including the primates (Jack & Campos, 2012; Peck et al., 2010) cats (Zapata-Ríos & Araguillin, 2013), and bears (Castellanos, 2011), as well as the frogs (Arteaga et al., 2016; Tapia et al., 2017), birds (Jahn, 2008; Willig & Presley, 2016) and orchids (Endara et al. 2009). **We recommend that the entire Bosque Protector system be extended the same protections as the SNAP system, particularly with regards to prohibition of mining.**

As water resources throughout the world increasingly come under pressure, unlogged watersheds in Bosques Protectores and other reserves are accordingly precious. The tropical montane cloud forests of Ecuador are particularly important for water cycling across a much larger area than they cover due to water capture by their biodiverse epiphytes, plants such as orchids that live on top of other plants. The epiphytes comb water out of the fog, helping these forests to capture up to 75% additional water through fog drip (Bruijnzeel et al., 2011; Cavelier, Solis, & Jaramillo, 1996), enabling cloud forests to maintain dependable flow downstream during dry periods (Bubb, May, Miles, & Sayer, 2004). Our results underscore that these montane forested ecosystems are valuable for not only for water, they also contain a very large number of rare species. Mining, particularly of copper and gold, will not only destroy the biodiversity and its water generating and holding capacity, but also strongly decrease the quality of water downstream—where people, invertebrates, and fish depend on it—for generations, by changing acidity and releasing toxic compounds such as mercury and arsenic (Bundschuh et al., 2012; Leblanc, Morales, Borrego, & Elbaz-Poulichet, 2000; Oyarzun et al., 2006).

Preservation of the primary forests in Bosques Protectores would allow the current economic benefits of these reserves to grow. It would also enable future economies through ethical and ecologically-minded bioprospecting by Ecuadorian researchers, leading to long-term economic returns for the people of Ecuador and scientific and medical rewards for all of humanity (Cragg & Newman, 2013; Harvey, 2000; Mathur & Hoskins, 2017; Rafiq et al., 2017; Strobel & Daisy, 2003). For example, a recently described species found at Los Cedros, *Cuatresia physalana* (Orozco & Canal, 2011), is related to tomatoes and potatoes and thus may contain genetic materials valuable for agriculture. Furthermore, *Cuatresia* are known to contain anti-malarial compounds (Deharo et al., 1992; Krugliak, Deharo, & Shalmiev, 1995). It is not only plants that are a source of antimicrobials and other bioactive compounds, so are plant‐ and soil-associated microbes (Cragg & Newman, 2013; Strobel & Daisy, 2003); microbes too are lost with deforestation and land conversion (Rodrigues et al., 2013).

In 2008, Ecuador set a new moral standard for the world when the National Assembly included the rights of Nature in the Constitution of Ecuador (articles 71–74, (Asamblea Nacional, 2008)). It is time to follow through on this commitment.

## Funding

The authors received no financial support for the research, authorship, and/or publication of this article.

## Acknowledgements

We are grateful to all the reserve managers, but especially José Decoux and Carlos Zorilla, for being on the frontlines. Carlos Morozch from Mashpi Lodge and Alejandro Solano, Agustina Arcos and Oliver Torres from Mashpishungo and Pambiliño preserves in Mashpi were helpful during stream surveys. Mashpi’s blog posts about nature were particularly informative about what species are there. We thank John Seed and the Rainforest Information Centre for their leadership in conservation and for the many conversations we have had over the last year. Danny Newman’s comments improved the manuscript. Shuheng Ni and Ali Luddon helped with the orchid table. Andreas Kaye, Michael Wherley, and Sidney Glassman provided access to photographs. Bruce Holst at Marie Selby Botanical Gardens and Susan Leon-Yanez at the QCA Herbarium provided plant voucher information. Inspiration for writing this paper came from checking a list of “potential” species (based on range maps) in the Intag by students at Cornell University for DECOIN (Defensa y Conservacion Ecologica de Intag). We wanted to know what was known to be present.

